# Mapping potential malaria vector larval habitats for larval source management: Introduction to multi-model ensembling approaches

**DOI:** 10.1101/2022.10.06.511086

**Authors:** Guofa Zhou, Ming-Chieh Lee, Xiaoming Wang, Daibin Zhong, Guiyun Yan

## Abstract

Mosquito larval source management (LSM) is a viable supplement to the currently implemented first-line malaria control tools for use under certain conditions for malaria control and elimination. Implementation of larval source management requires a carefully designed strategy and effective planning. Identification and mapping of larval sources is a prerequisite. Ensemble modeling is increasingly used for prediction modeling, but it lacks standard procedures. We proposed a detailed framework to predict potential malaria vector larval habitats using ensemble modeling, which includes selection of models, ensembling method and predictors; evaluation of variable importance; prediction of potential larval habitats; and assessment of prediction uncertainty. The models were built and validated based on multi-site, multi-year field observations and climatic/environmental variables. Model performance was tested using independent multi-site, multi-year field observations. Overall, we found that the ensembled model predicted larval habitats with about 20% more accuracy than the average of the individual models ensembled. Key larval habitat predictors were elevation, geomorphon class, and precipitation 2 months prior. Mapped distributions of potential malaria vector larval habitats showed different prediction errors in different ecological settings. This is the first study to provide a detailed framework for the process of multi-model ensemble modeling. Mapping of potential habitats will be helpful in LSM planning.

**Author’s summary:** Mosquito larval source management (LSM) is a viable supplement to the currently implemented first-line malaria control tools. Implementation of LSM requires a carefully designed strategy and effective planning. Identification and mapping of larval sources is a prerequisite. Ensemble modeling is increasingly used for prediction modeling, but it lacks standard procedures. We proposed a detailed framework for such a process, including selection of models, ensembling methods and predictors; evaluation of variable importance; and assessment of prediction uncertainty. We used predictions of potential malaria vector larval habitats as an example to demonstrate how the procedure works, specifically, we used multi-site multi-year field observations to build and validate the model, and model performance was further tested using independent multi-site multi-year field observations – this training-validation-testing is often missing from previous studies. The proposed ensemble modeling procedure provides a framework for similar biological studies.

## Introduction

Malaria is still the most serious mosquito-borne infectious disease in the tropics, especially in Africa [1]. The scale-up of indoor interventions such as long-lasting insecticidal nets (LLINs) and indoor residual insecticide spraying (IRS), together with effective treatment, has led to a substantial reduction in the malaria burden [1]. Nonetheless, malaria control faces increased challenges due to vector resistance to insecticides and outdoor residual transmission [1,2]. Larval source management (LSM) has become a viable choice for further reducing malaria transmission and is recommended by the World Health Organization (WHO) for use under certain conditions for malaria control and elimination [3,4]. Larval control complements LLINs and IRS and controls both indoor and outdoor transmission [3,5,6]. Previous studies in several malaria-endemic countries prove that larval source reduction and larviciding can significantly reduce both indoor and outdoor vector density and malaria infections [5,7–10]. Several African countries have adopted LSM as a key vector control tool parallel to LLINs and IRS or as a supplementary strategy [11–13]. For example, the Kenyan government has planned to target all larval sources for LSM by 2023, although the plan is over-ambitious [11]. Because larval sources can be anywhere after the rain, the currently recommended LSM strategy is targeted LSM with environmental management and larviciding [3]. Large-scale implementation of LSM requires a carefully designed strategy and effective planning, especially the identification and mapping of larval sources prior to any field operations [11–14]. However, effective larval habitat identification and mapping is lacking.

Many climatic- and environmental-based models have been used to predict the potential distribution of malaria vector larval habitats, for example, ecological/environmental niche models [15,16], logistic regression [17–19], and machine learning methods such as artificial neural network and random forest models [17,18]. Clearly, different modeling methods are likely to produce different results and select different risk factors [17,18,20]. With so many models available, it will be difficult to find the robust model with the best predictions. More importantly, since different models end up with different groups of risk factors, how should the key risk factors be determined?

Recently, multi-model ensemble approaches have increasingly been used for predictions in various field studies [21–28]. According to Kotu and Deshpande, “Ensemble modeling is a process where multiple diverse models are created to predict an outcome, either by using many different modeling algorithms or using different training data sets. The ensemble model then aggregates the prediction of each base model and results in one final prediction for the unseen data” [29]. The reasons for employing ensemble methods in building a model are to enhance the overall performance of the model, minimize the error rate that can be caused by using individual models, and reduce the overall uncertainty of predictions [22,29,30]. There are different ways to ensemble the models, including most-votes, simple average, weighted average (linear or nonlinear), boosting, and stacking [22,24,27,31–33]. In mosquito studies, ensemble modeling has been used to predict the global expansion of *Aedes* mosquitoes and the invasion of *Anopheles stephensi* in Africa [26,34,35]. Very recently, ensembled modeling techniques have also been adopted to predict potential mosquito larval habitats [20,25,36]. For example, Wieland et al. used the average of dry/wet/normal year predictions as the ensembled prediction, which biologically makes sense for reducing over-or underestimation due to variations in precipitation [36], although we don’t know whether the dry and wet years have similar intensity (negative/positive) of impact on larval habitat availability and productivity. In studies by Rhodes et al. and Beeman et al., the final ensembled model was fitted using only those models with area-under-curve (AUC) scores of ≥0.7, and each model was weighted proportionally to its AUC score [20,25]. The assumption is that only models with AUC > 0.7 are considered to be well-performing models, and better-performing models should be given higher weight [20,24,25,27]. These methods lack biological (and/or statistical) basis for both the model and weight selections; the selection of AUC > 0.7 is somewhat arbitrary and the weight selection may not lead to robust estimations. More importantly, feature importance or risk factor analysis is essential for larval habitat prediction and for LSM [21,36]. Bose et al. used rank-order (i.e., most votes) for feature selection [21]. Rhodes et al. and Sinka et al. did not specify how the variables were selected for the ensembled model [25,26]. Beeman et al. displayed the variable importance for each individual model but not for the ensembled model [20]. Overall, no standard method currently exists for variable selection and relative importance evaluation for ensembled models [24,27,29,37].

The aim of this study was to build a habitat prediction model using historical observations of aquatic and larval positive habitats in western Kenya and weighted multi-model ensembling. We used recent field observations and observations from different areas to test the model. Finally, we proposed a method to assess risk factors and to measure the uncertainty of model predictions. The predicted map of habitat and larval distribution in western Kenya will be beneficial for LSM planning.

## Materials and methods

### Study area, field data collection and data assignment for modeling

Field aquatic habitats and mosquito larval surveys were conducted in six sentinel sites in Kakamega (Iguhu site), Vihiga (Mbale, Emakakha and Emutete sites), Kisumu (Kombewa site) and Kisii (Marani site) counties in western Kenya (Figure 1). Western Kenya is the last malaria transmission hotspot persisting in Kenya [38–43]. The study sites included places with seasonal (e.g., Marani) and perennial (e.g., Kombewa) malaria transmission. Malaria larval distribution, larval ecology and parasite transmission in these sites have been extensively studied in the past 20 years [38–52]. Three sites, Kombewa, Emakakha and Emutete, are relatively flat with shallow valleys. Kombewa is located in the Lake Victoria shore area with an elevation range from 1,150m to 1,350m above sea level (a.s.l.), and Emakakah and Emutete have an elevation range from 1,420m to 1,520m a.s.l. Iguhu and Mbale are hilly areas with elevation from 1,420m to 1,670m, and Marani has an elevation of 1,520–1,760m a.s.l. with steep valleys. The study area has two rainy seasons: a long rainy season that usually starts in April and lasts until July, and a short rainy season between October and November, with two dry seasons in between [43].

**Figure 1.**
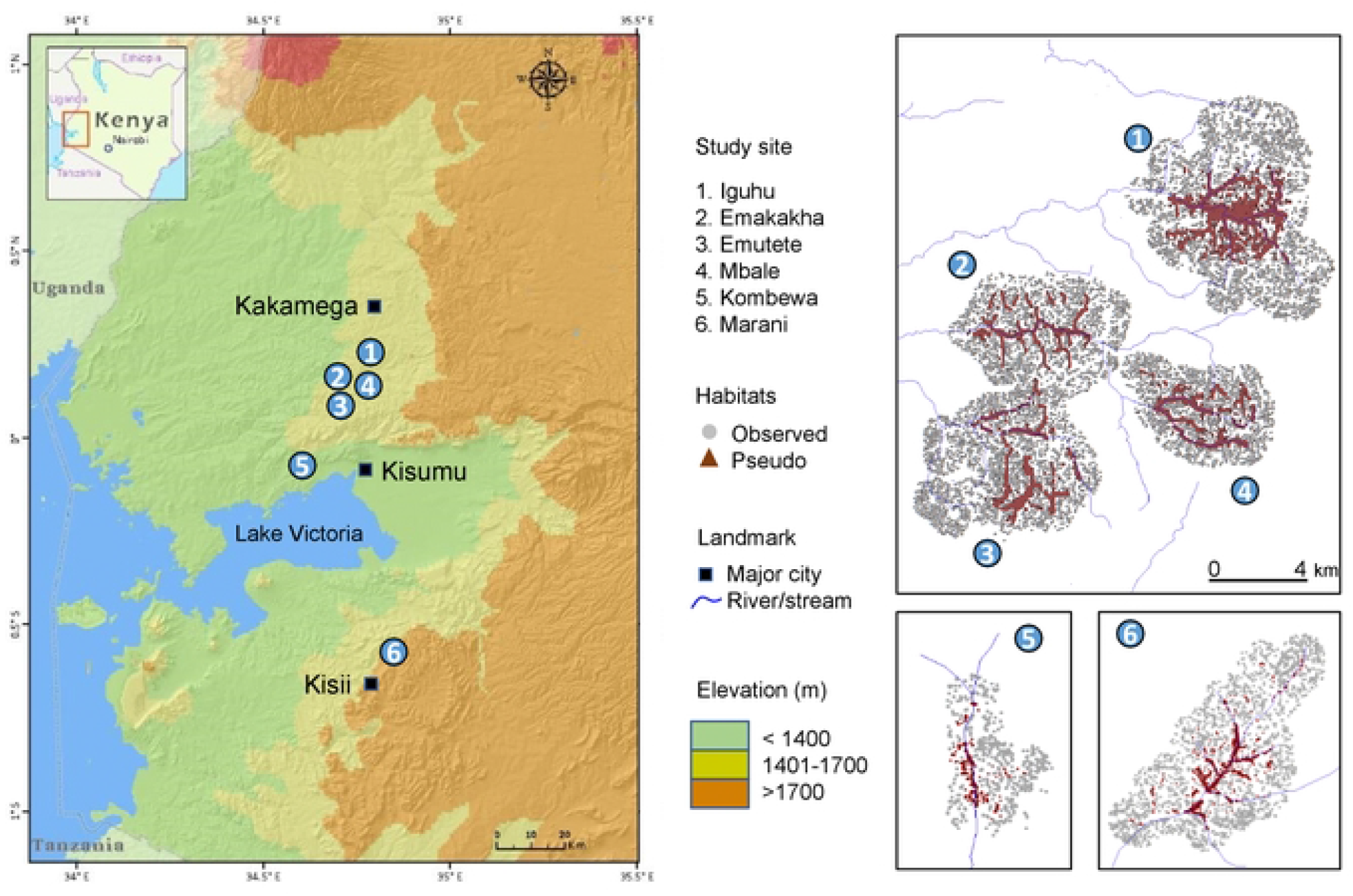
Study sites (left) and distribution of observed aquatic habitats (2003–2018) and pseudo-habitats in the six study sites in western Kenya

Annual precipitation ranges from around 1,400mm in Kakamega, Vihiga and Kisumu counties to about 1,700mm in Kisii County. Aquatic habitats and mosquito larval infestations were surveyed in 2001, 2003, 2005, 2008, 2010–2012, 2017 and 2018 (Figure 1). In most years, field habitat surveys were carried out in February, May, August and November. GPS locations of all habitats were recorded, sizes of habitats were measured, and availability (yes/no) of immature *Anopheles* mosquitoes was checked in most years. Overall, we surveyed about 50,000 aquatic habitats, and *Anopheles* larval infestation status was available in about 40,000 habitats (Table 1). We have noticed that very few studies have predicted the potential of larval positive habitats [19].

**Table 1.**
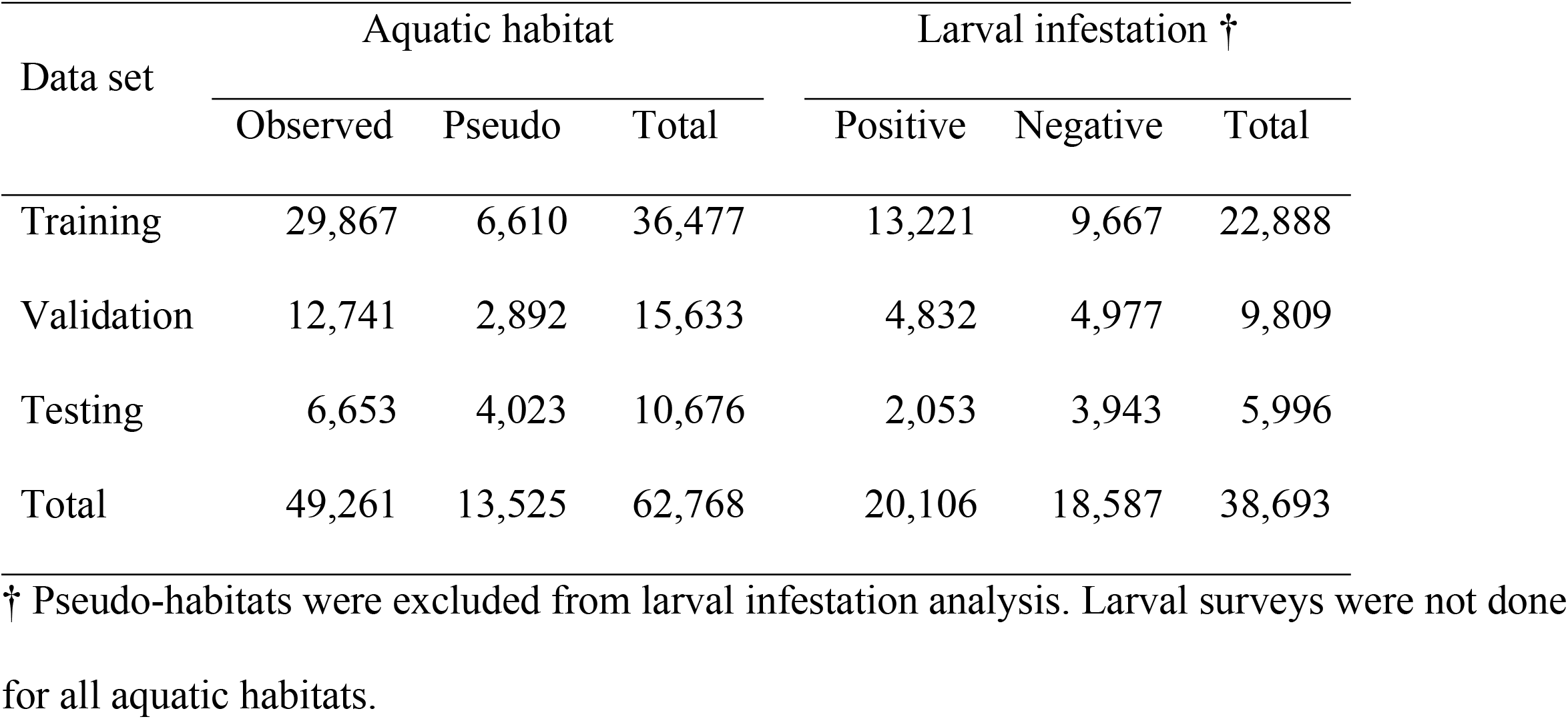
Number of samples used for model training, validation and testing for the identification of aquatic habitats and larval infested habitats

Since all aquatic habitats had water at the time they were surveyed, to identify predictors of aquatic habitats we artificially generated about 13,500 pseudo-habitats in randomly selected locations with known residential houses [25,36] (Table 1). Pseudo-habitats were selected at least 50m away from any known aquatic habitats. These pseudo-habitats will always have aquatic status as ‘no water’ regardless of sampling season. The pseudo-habitats were generated using ArcGIS 10.0 (ESRI, Redlands, California 92373, USA).

We used 2003–2012 field observations from the Iguhu, Emakakha, Emutete and Mbale sites to train and validate the models. A 70:30 random splitting of data was used for model training and validation (Table 1). To produce an unbiased evaluation of the final model, we used field data collected from the same area in 2017 as testing data, i.e., an independent data set [53,54]. To further independently evaluate model performance, we used data collected from the Kombewa and Marani sites for model testing. About two-thirds of the pseudo-habitats were randomly assigned (with equal probability) into February, May, August and November 2012 samples, and one-third were assigned as 2017 samples.

### Climatic and environmental data

Climatic and environmental data included about 220 variables and has been described in a previous study [43]. Briefly, environmental data included DEM, topographic/geomorphologic features (e.g., slope, aspect, land surface roughness, etc.), land use land cover, and tree coverages from different years. Climatic data included monthly average temperature and cumulative precipitation. Satellite image–derived data included variables such as normalized difference vegetation index (NDVI), normalized difference water index (NDWI), land surface temperature, evapotranspiration, etc. For monthly data, we selected only the data from the six months prior to the habitat surveys, assuming habitat aquatic status and larval availability were affected only by climatic conditions during the last six months.

The assignments of climatic and environmental data to each survey point were done using ArcGIS 10.0.

### Modeling process

Slightly different from the single-model modeling process [43], ensemble modeling requires an extra step to build all individual models a priori (Figure S1). Data dimension reduction was done using principal component analysis (PCA) and has been described in previous studies [29,43]. Individual model training and prediction are rather straightforward; however, there is no standard method for model ensembling or risk factor determination in ensemble modeling [20,25,26,29,30,36,55].

### Model specification and modeling process

Many models can be used to identify mosquito larval habitats [17,18,20,25]. There is no standard method for model selection in ensemble modeling. Many studies have used the R package *biomod2* for model selection and ensembling [20,25,26]. However, in principle, model diversity and independence are key to ensemble modeling [29,37], because models built using similar methods may perform similarly (for example, gradient-boosted logistic and gradient-boosted tree models), so ensembling them may provide limited additional information for risk factor analysis and limited improvement in model performance due to redundancy. Including conventional models such as logistic and decision tree models and modern machine learning methods such as neural network increases model diversity. We selected 10 typical models for classification analyses in this study; five of them are conventional methods and five are machine learning methods (Table 2).

**Table 2.** Models used in this study

| Type of model | Specific model | Conventional vs.<br>machine learning |
| --- | --- | --- |
| Logistic regression | Ordinary logistic | Conventional |
|  | Bayesian logistic | Conventional |
|  | Gradient-boosted (GB) logistic | Machine learning |
|  | Support vector machine (SVM) logistic | Machine learning |
| Decision tree | Regular classification tree | Conventional |
|  | Naïve Bayes tree | Conventional |
|  | k-nearest neighbor (kNN) classifier | Conventional |
|  | Extreme gradient-boosting (XGB) tree | Machine learning |
|  | Random forest | Machine learning |
| Neural network | Regular neural network tree | Machine learning |

Two of the 10 models require careful prior specifications: Bayesian logistic regression and neural network (NNW). For Bayesian logistic regression, one needs to specify a joint distribution for the outcome(s) and all the unknown parameters, which typically takes the form of a marginal prior distribution for the unknowns multiplied by a likelihood for the outcome(s) conditional on the unknowns. In this study, since we had 0-1 (yes-no) outcomes, we used a conditional binomial model with a logit link function. Since we had little a priori confidence that the parameters would be close to zero, we chose to use Student’s *t*-distribution as the prior distribution for parameter estimation, i.e., heavier tails than the normal-like shape. For the NNW model, we used a logistic activation function for the output layer and tangens hyperbolicus for the hidden layer(s). We also tested different hidden layer structures for the NNW, ranging from 1 to 3 layers and 2 to 5 neurons for each layer. All models need some prior specifications, but most of the settings are straightforward and likely do not severely affect the classification results, for example, the number of repeats of cross-validations (10 in this study) and the number of trees (as long as we specify a large number).

We used stepwise variable selection in ordinary logistic regression and projection predictive variable selection in Bayesian logistic regression [56,57]. Variable selection for other models was based on either a significance test (P < 0.05) or relative importance score (<1%) [29]. In addition to using a separate data set for validation, to further reduce predictive bias and uncertainty (i.e., variance of performance estimates) we used 10-fold cross-validation (10-fold cv) for the training of all models except NNW, since we had a large training data set [29,53,58,59].

For aquatic habitat identification, we used both field-observed aquatic habitats (yes) and pseudo-habitats (no). For the identification of larval-positive habitats, we excluded the pseudo-habitats.

### Comparisons of model performance

We used several criteria to evaluate the performance of different models. The overall prediction accuracy was calculated for each model using both the validation and testing data sets. Validation was done using the original model built on the training data set. Prediction accuracy of the final model was assessed using the testing data set, and the model was calibrated using the validation data set. Since we had binary outcomes, we also calculated the sensitivity, specificity, area under curve (AUC), positive/negative likelihood ratios and positive/negative predictive values for each model [37,59,60]. Agreement between observed and predicted records was measured using the Kappa statistic [61,62].

To assess other differences in model performance, we checked the relative importance of variables selected for model predictions. Variable relative importance was measured using scaled relative importance ranging from 0 (least important) to 100 (most important).

### Ensembling of models

Since different models may select different variables for predictions (i.e., harness different aspects of the data) the models likely don’t contribute equally to the ensembled model. To obtain robust predictions, the ensembling process can be treated as a classic case of simplex optimization. Assuming *p*_i_ is the output from each model and *w*_i_ is the weight for each model such that 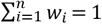, where *n* is the total number of models, the predicted value using the ensembled model is

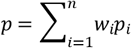

The objective function is a maximum likelihood function (MLF), i.e.,

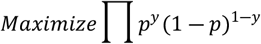

where *y* is the response flag for the observations. This is a classic logistic regression type of MLF. Thus, the weights *w*_i_ can be estimated using logistic regression analysis. We also examined the weight estimates by using the simple average (equal weight for all models), the most votes (predictions by the most models) and neural network models (to measure potential nonlinearity).

The ensembled model used predictions based on all data sets; i.e., models were calibrated by all data sets rather than the training data set alone.

### Ensembling of risk factors

Currently there is no standard method for examining variable importance in ensembled models [20,25,29,60]. We proposed the following approach. For each model, the top 20 most important (by relative influence) risk factors (predictors) were selected. For any model (e.g., logistic regression) that ended up with <20 significant risk factors after variable selection, only the significant risk factors were selected. The risk factors were ranked based on the votes of the models (i.e., how many models selected each risk factor), and the top 20 ranked risk factors were selected as the important risk factors. The variable importance was measured as the weighted average of the relative influence from each individual model:

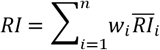

where 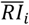 was the standardized (scaled) relative importance *RI*_i_ from model *i*, i.e.,

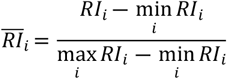

The weights, *w*_i_, were the same as estimated by the ensembled model.

### Prediction uncertainty

Model prediction uncertainty has also less studied or not mentioned in most previous studies. We proposed to use the square-rooted mean-squared error (MSE) against the ensembled model predictions to measure the prediction uncertainty [29,37], i.e.,

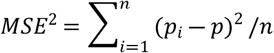

where *p* was the probability predicted by the ensembled model and *p*_i_ was the probability predicted by each individual model.

All data analyses were conducted using R 4.0.3 (The R Foundation for Statistical Computing, Vienna, Austria). The following packages were used in this study: *caret* for training and validation data splitting and for kNN tree; *factoextra, Rcpp, FactoMineR, ggplot2, Hmisc* and *reshape2* for data dimension reduction; *dplyr, ROCR, caTools, mlbench, MLmetrics, MASS, plyr* and *tidyverse* for logistic regression; *mboost, gbm* and *cvAUC* for GBM logistic; *arm, logicFS, LogicREG* and *mcbiopi* for SVM logistic; *GGally, bayesplot, rstanarm, loo, projpred* and *reportROC* for Bayesian logistic; *party, rpart* and *pROC* for decision tree; *randomForest* for random forest; *Metrics* and *xgboost* for XGB tree; *neuralnet, devtools* and *usethis* for NN models; *deepnet* for DeepNN models, *e1071* for Naïve Bayes classifier, and *vip* for measuring variable importance. The above packages are listed in the order they appeared in the code. Some packages were used for multiple models; for example, *caret* was used for data partitioning and in logistic regression, SVM logistic, GBM tree, and XGB tree, among others.

## Results

This study collected data from 49,261 aquatic habitats, of which 38,693 had larval survey results (Table 1). For aquatic habitat identification, about 42,600 aquatic habitat records were used for model training and validation and 6,600 for model testing. For larval positive habitat identification, about 32,000 records were used for model training and validation and 6,000 for model testing.

### Prediction accuracy

For aquatic habitat identification, prediction accuracy for the testing data varied substantially among different models (Figure 2A); the overall prediction accuracy varied from 58% (neural network) to 82% (ensembled model) (Table 3). For predicting the habitats in Kakamega and Vihiga counties, the ensembled model (accuracy 89.7%) did not produce the best accuracy; however, for Kisii (accuracy 78.1%) and Kisumu (accuracy 68.2%) counties and overall (accuracy 82.3%), the ensembled model was the best choice (Figure 2A). For the overall prediction, logistic regression ensembling outperformed simple average, most-votes, and neural network–ensembled models (Supplementary Figure S2A).

**Table 3.** Evaluation of model performance for predicting the aquatic habitats using testing data set from the six study sites

| Evaluation indicator | Regular<br>logistic | Bayesian<br>logistic | GBM<br>logistic | SVM<br>logistic | Decision<br>tree | Naïve<br>Bayes | k-NN<br>tree | XGB<br>tree | Random<br>forest | Neural<br>network | Ensembled |
| --- | --- | --- | --- | --- | --- | --- | --- | --- | --- | --- | --- |
| Kappa agreement | 0.30 | 0.53 | 0.25 | 0.35 | 0.27 | 0.32 | 0.42 | 0.00 | 0.32 | 0.22 | 0.62 |
| **Accuracy rate** | **0.64** | **0.77** | **0.60** | **0.65** | **0.58** | **0.66** | **0.71** | **0.62** | **0.62** | **0.58** | **0.82** |
| Area under curve (AUC) | 0.68 | 0.68 | 0.78 | 0.73 | 0.84 | 0.65 | 0.72 | 0.82 | 0.65 | 0.50 | 0.80 |
| Sensitivity | 0.57 | 0.78 | 0.46 | 0.50 | 0.33 | 0.65 | 0.70 | 1.00 | 0.40 | 0.43 | 0.88 |
| Specificity | 0.76 | 0.76 | 0.83 | 0.89 | 0.99 | 0.68 | 0.74 | 0.00 | 0.98 | 0.83 | 0.73 |
| Positive predicted value | 0.80 | 0.84 | 0.82 | 0.88 | 0.99 | 0.77 | 0.81 | 0.62 | 0.98 | 0.80 | 0.84 |
| Negative predicted value | 0.52 | 0.68 | 0.48 | 0.52 | 0.47 | 0.54 | 0.60 | 1.00 | 0.50 | 0.47 | 0.79 |
| Positive likelihood ratio | 2.37 | 3.20 | 2.69 | 4.62 | 55.78 | 2.06 | 2.66 | 1.00 | 24.76 | 2.49 | 3.23 |
| Negative likelihood ratio | 0.57 | 0.29 | 0.65 | 0.56 | 0.67 | 0.51 | 0.41 | 0.00 | 0.61 | 0.69 | 0.16 |

**Figure 2.**
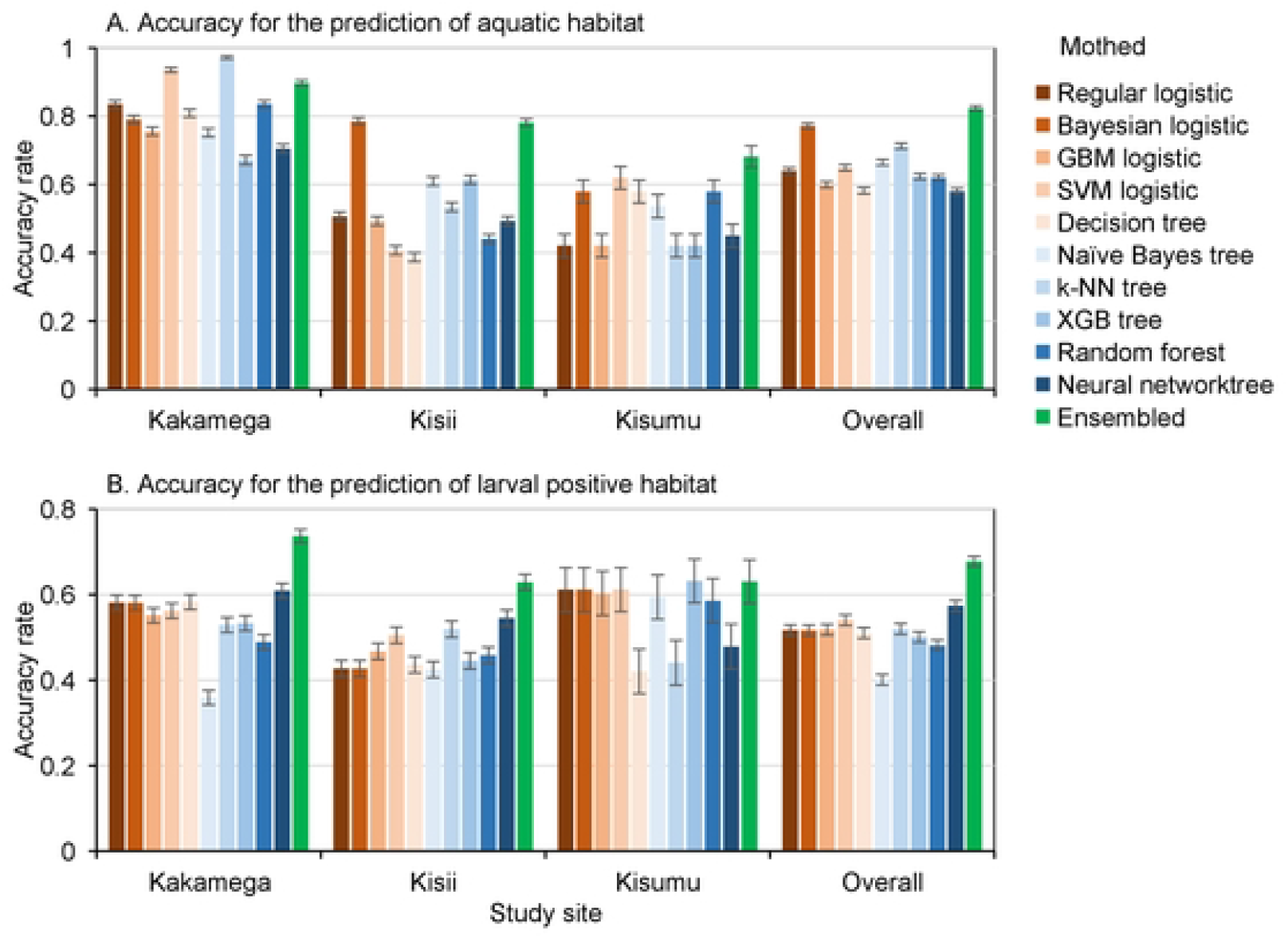
Accuracy of predictions of different models at different sites. A: Prediction of aquatic habitats; B: Prediction of larval positive habitats.

Similarly, for larval infestation identification, prediction accuracy varied substantially among different models at different sites (Figure 2B), and all models had <60% prediction accuracy for the testing data except the ensembled model (accuracy 68%) (Table 4). In nearly all counties and overall, the ensembled model overperformed other models (Figure 2B), although prediction accuracy was lower compared to the aquatic habitat identification models (Figure 2A). Also similarly, logistic regression ensembling overperformed simple average, most-votes and neural network ensembled models (Supplementary Figure S2B).

**Table 4.**
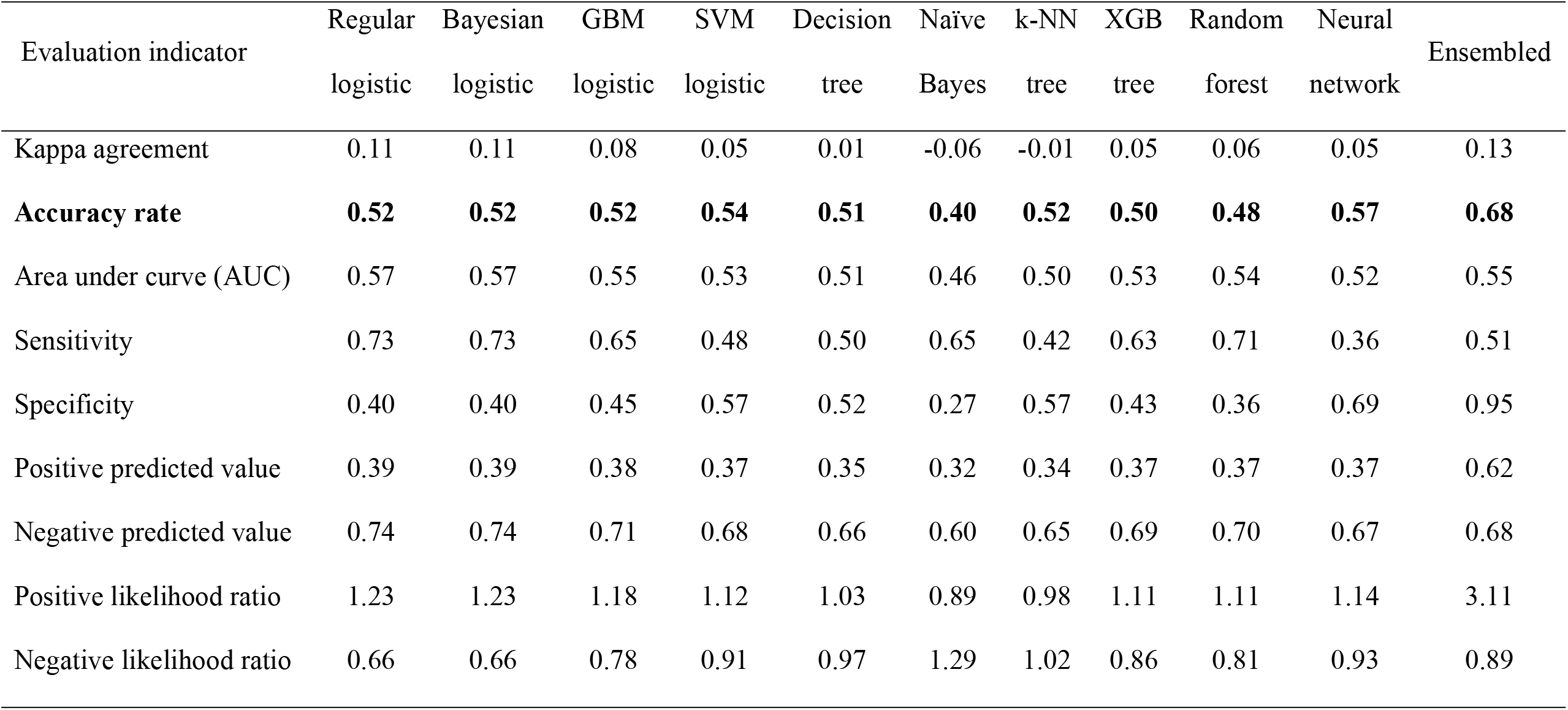
Evaluation of model performance for the prediction of larval positive habitats using testing data set from the six study sites

On average, the logistic regression–ensembled model had about 18% greater accuracy than each individual model for the prediction of aquatic and larval positive habitats (Figure 2, Tables 3 & 4).

### Agreement between observations and predictions

For the aquatic habitat identification, prediction sensitivity of the 10 models ranged from 33% to 100% and specificity ranged from 0% to 99%, and the overall Kappa agreement was low (<0.6) for all models (Table 3). The ensembled model had a Kappa of 0.62 (i.e., moderate agreement), 88% sensitivity, 73% specificity, and 80% AUC – reasonably good accuracy compared to pure chance (AUC of 0.5) (Table 3). The ensembled model had about 30% higher sensitivity than the average of individual models.

For the larval positive habitats, all models had relatively low sensitivity and specificity, and all AUC < 60% (Table 4). Kappa agreement between observed and predicted was poor (<0.2 for all). The ensembled model had similar sensitivity to the average of individual models (∼50%), but the ensembled model had 50% higher specificity compared to the average of individual models (Table 4).

For the final model, aquatic habitat prediction had an AUC of 97.9%, Kappa agreement 0.81 (95% CI [0.805, 0.822]), sensitivity of 95.6% and specificity of 85.0%. For the larval positive habitat prediction, the AUC was 74.1%, Kappa agreement 0.35 (95% CI [0.334, 0.365]), sensitivity 50.4% and specificity 83.2%.

### Identification of risk factors

It is clear that different models, in most cases, selected different groups of important risk factors because of the difference in variable selection algorithms (Tables S1 & S2). In some models, a few risk factors had clear high relative influences, for example, the GBM logistic and XGB tree models for identifying aquatic habitats (Table S1). In other models, many factors influenced the model predictions, for example, random forest for the prediction of larval positive habitats (Table S2).

If we take the ensembled model as the final model, DEM (relative influence 100%), geomorphon class (rel.inf 72.9%) and the amount of precipitation 2 months prior to the survey (rel.inf 45.9%) were the top three risk factors determining the aquatic habitats (Table 5). The relative influence of the other factors was significantly lower compared to the top three (Table 5). While many factors may have affected larval infestation (Table 6), the top three risk factors were maximum temperature 4 months prior to the survey, the amount of precipitation 3 months prior to the survey, and distance to the river/streams; however, northness and other factors also had high influence (Table 6). Regardless, the top 10 risk factors were all related to temperature and precipitation with the exception of northness, which is somewhat related to the amount of sunlight received (Table 6).

**Table 5.**
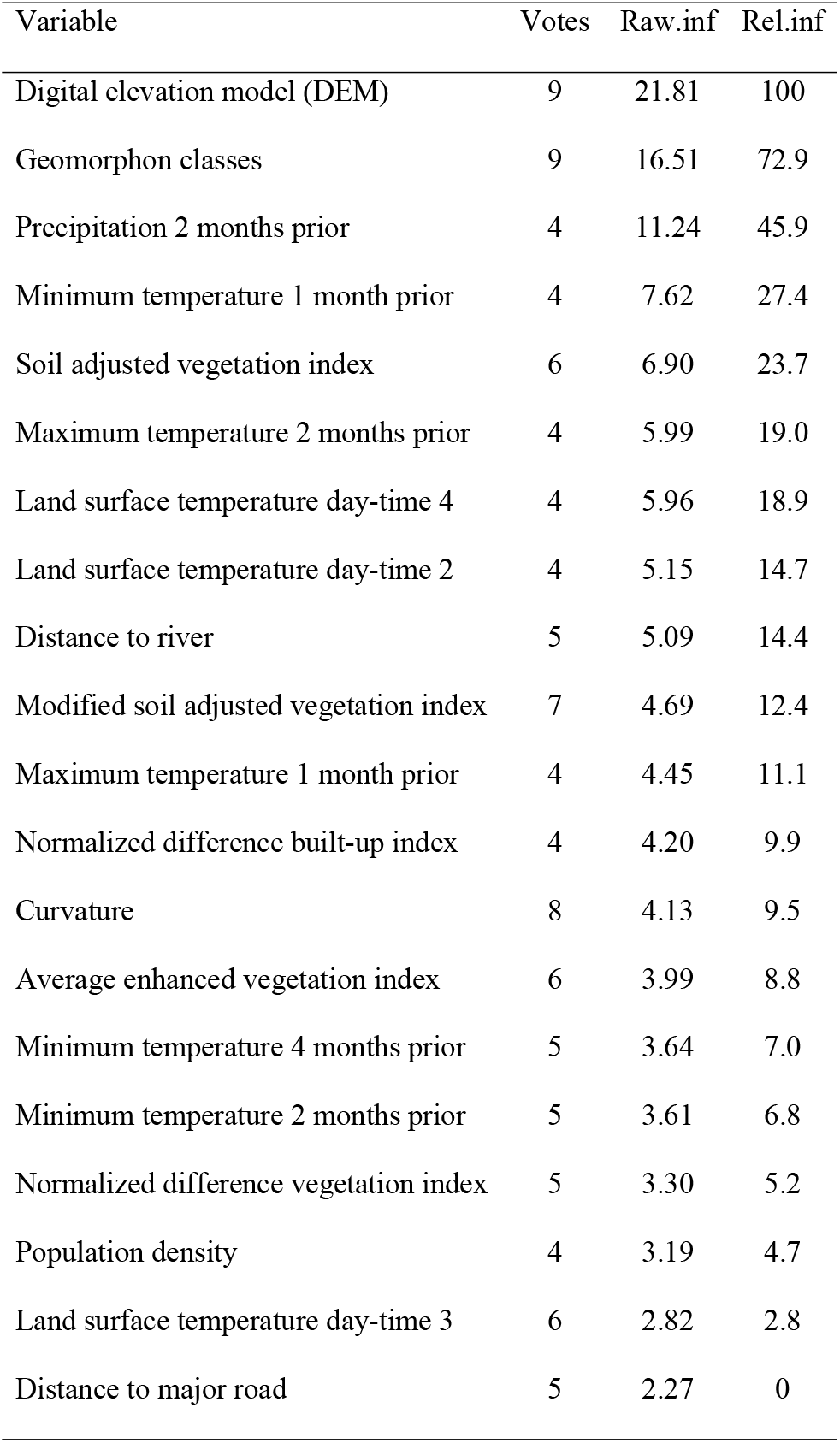
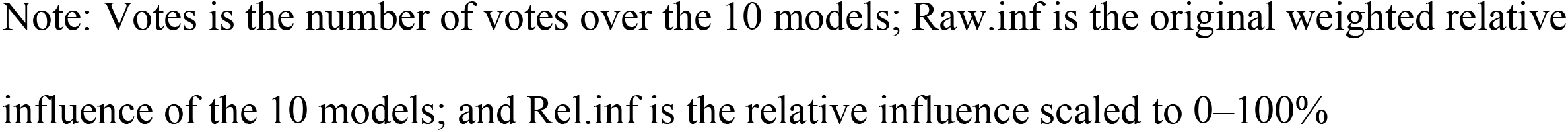
Relative influence of the top 20 risk factors for the prediction of aquatic habitats

| Variable | Votes | Raw.inf | Rel.inf |
| --- | --- | --- | --- |
| Digital elevation model (DEM) | 9 | 21.81 | 100 |
| Geomorphon classes | 9 | 16.51 | 72.9 |
| Precipitation 2 months prior | 4 | 11.24 | 45.9 |
| Minimum temperature 1 month prior | 4 | 7.62 | 27.4 |
| Soil adjusted vegetation index | 6 | 6.90 | 23.7 |
| Maximum temperature 2 months prior | 4 | 5.99 | 19.0 |
| Land surface temperature day-time 4 | 4 | 5.96 | 18.9 |
| Land surface temperature day-time 2 | 4 | 5.15 | 14.7 |
| Distance to river | 5 | 5.09 | 14.4 |
| Modified soil adjusted vegetation index | 7 | 4.69 | 12.4 |
| Maximum temperature 1 month prior | 4 | 4.45 | 11.1 |
| Normalized difference built-up index | 4 | 4.20 | 9.9 |
| Curvature | 8 | 4.13 | 9.5 |
| Average enhanced vegetation index | 6 | 3.99 | 8.8 |
| Minimum temperature 4 months prior | 5 | 3.64 | 7.0 |
| Minimum temperature 2 months prior | 5 | 3.61 | 6.8 |
| Normalized difference vegetation index | 5 | 3.30 | 5.2 |
| Population density | 4 | 3.19 | 4.7 |
| Land surface temperature day-time 3 | 6 | 2.82 | 2.8 |
| Distance to major road | 5 | 2.27 | 0 |

Note: Votes is the number of votes over the 10 models; Raw.inf is the original weighted relative influence of the 10 models; and Rel.inf is the relative influence scaled to 0–100%

**Table 6.**
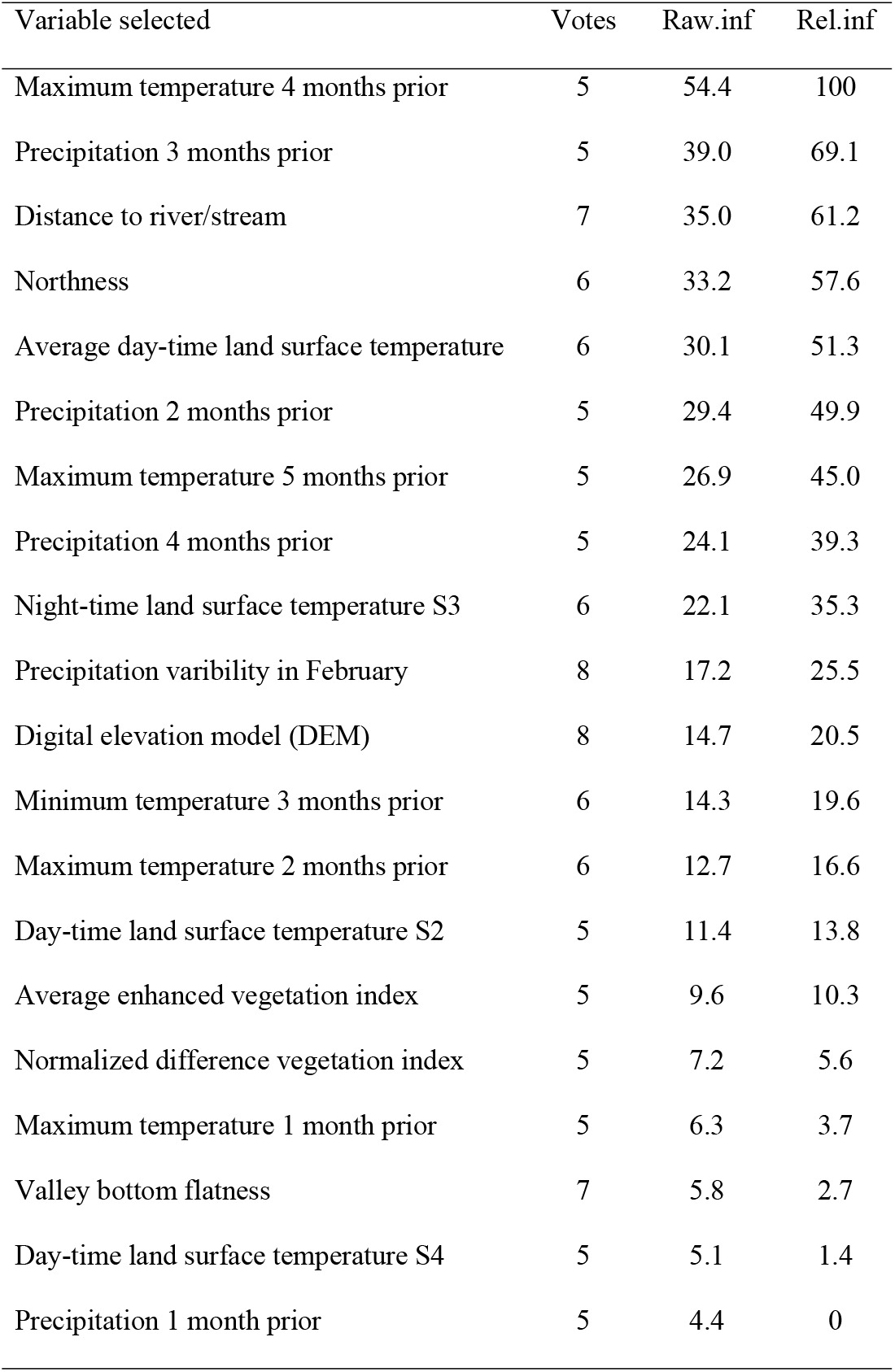
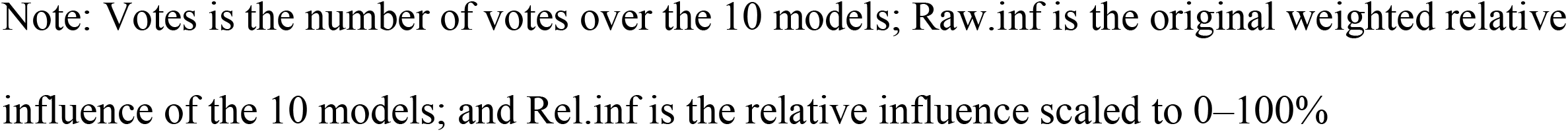
Relative influence of the top 20 risk factors for the prediction of larval positive habitats

| Variable selected | Votes | Raw.inf | Rel.inf |
| --- | --- | --- | --- |
| Maximum temperature 4 months prior | 5 | 54.4 | 100 |
| Precipitation 3 months prior | 5 | 39.0 | 69.1 |
| Distance to river/stream | 7 | 35.0 | 61.2 |
| Northness | 6 | 33.2 | 57.6 |
| Average day-time land surface temperature | 6 | 30.1 | 51.3 |
| Precipitation 2 months prior | 5 | 29.4 | 49.9 |
| Maximum temperature 5 months prior | 5 | 26.9 | 45.0 |
| Precipitation 4 months prior | 5 | 24.1 | 39.3 |
| Night-time land surface temperature S3 | 6 | 22.1 | 35.3 |
| Precipitation variability in February | 8 | 17.2 | 25.5 |
| Digital elevation model (DEM) | 8 | 14.7 | 20.5 |
| Minimum temperature 3 months prior | 6 | 14.3 | 19.6 |
| Maximum temperature 2 months prior | 6 | 12.7 | 16.6 |
| Day-time land surface temperature S2 | 5 | 11.4 | 13.8 |
| Average enhanced vegetation index | 5 | 9.6 | 10.3 |
| Normalized difference vegetation index | 5 | 7.2 | 5.6 |
| Maximum temperature 1 month prior | 5 | 6.3 | 3.7 |
| Valley bottom flatness | 7 | 5.8 | 2.7 |
| Day-time land surface temperature S4 | 5 | 5.1 | 1.4 |
| Precipitation 1 month prior | 5 | 4.4 | 0 |

Note: Votes is the number of votes over the 10 models; Raw.inf is the original weighted relative influence of the 10 models; and Rel.inf is the relative influence scaled to 0–100%

### Mapping potential larval habitats and uncertainty assessment

Risk mapping is rather straightforward. Figures 3 & 4 show the ensembled model predicted probability of potential aquatic habitats and larval positive habitats in western Kenya and the uncertainty of the predictions measured as mean-squared error. In both cases, predictions in Kakamega and Vihiga counties were more accurate in identifying aquatic and larval positive habitats (Figures 3 & 4). Predictions in Kisumu and Kisii counties had either lower probability (in Kisii) or higher error (Kisumu) compared to the other two counties (Figures 3 & 4).

**Figure 3.**
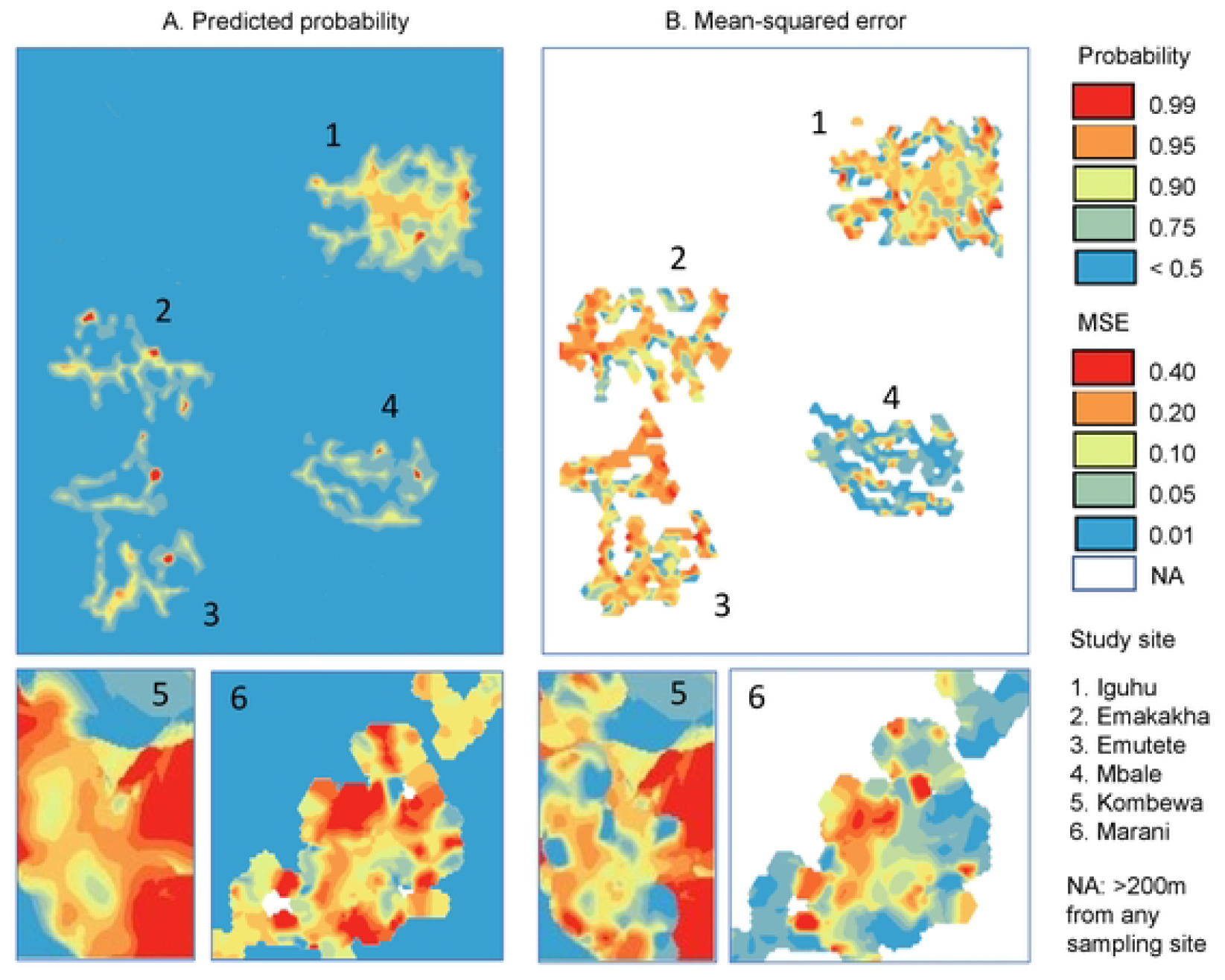
Predicted probability and mean-squared error of *Anopheles* larval positive habitats

**Figure 4.**
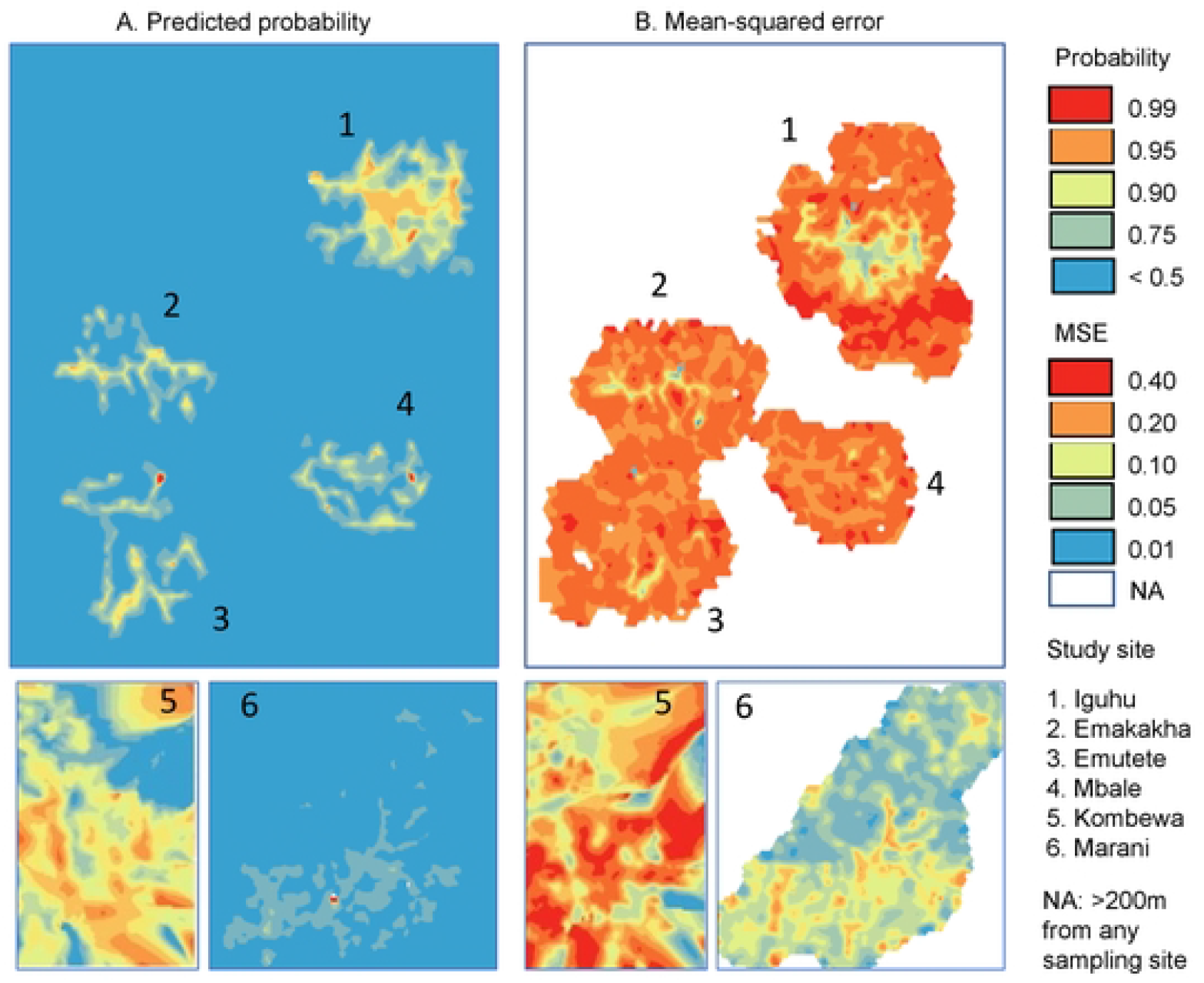
Predicted probability and mean-squared error of larval aquatic habitats

## Discussion

Larval source management is a viable supplement to the currently implemented first-line malaria control tools for use under certain conditions for malaria control and elimination [1,3,4]. The WHO has recommended LSM, and a number of African malaria-endemic countries have adopted LSM as a key vector control tool parallel to LLINs and IRS or as a supplementary strategy for malaria control and elimination [3,4,11–14]. Implementation of LSM requires a carefully designed strategy and effective planning; identification and mapping of larval sources is a prerequisite for LSM [11,12,14]. Using climatic/environmental data, especially satellite monitoring data, and mathematical models is a common approach for larval source identification [17–20,25,36]. Ensemble modeling provides high-accuracy larval source predictions; however, standard procedures are lacking, especially for ensembling method and risk factor selection and evaluation. Here we proposed a framework for larval source prediction using multi-model ensemble approaches, including methods for model selection, model ensembling, risk factor selection and evaluation, model prediction assessment, larval source mapping, and uncertainty analysis. To illustrate the procedure of the proposed approach, we used 10 years of multi-site field observations of larval habitat surveys for model training and validation, and multi-site multi-year (over seven years) independent field data for model testing. Using ensembles of 10 models, we identified three major predictors of aquatic habitat in western Kenya: elevation, geomorphon class, and amount of precipitation 2 months prior. We provided a map of potential malaria vector larval habitats in the study area. The aquatic habitat risk map will be valuable for LSM planning.

### Model selection in ensemble modeling

Data mining scientists recommend using diverse and independent models for ensemble modeling [29,37]. Most medical and biological application studies have not mentioned how individual models were selected for ensemble modeling [20,21,24–26,32,60]. However, model selection may affect the eventual predictions of the ensembled model. For example, in this study, we tested both GBM logistic and GBM classification models. The two models selected exactly the same group of variables with minimal difference in variable importance and very similar prediction results (results not shown). If we included both models in the ensembled model and used accuracy or AUC as the weight [24,25], we would likely over-weight (create bias toward) these models. Indeed, stepwise selection selected only the GBM logistic model (results not shown); i.e., the GBM classification model may be redundant. In fact, if we used accuracy or AUC > 0.7 for model selection in this study [24,25], we would actually end up with no model for larval positive habitat prediction. This is a reminder that the combination of artificially selected weights/models may severely affect the results of the ensembled model.

### Relative importance of risk factors

Currently there is no standard method in ensemble modeling for selecting the important predictors and estimating the relative influence of these predictors [29,37]. In cancer research, Liu et al. used the sum of relative importance across all models [60], which is equivalent to the simple average over all models, i.e., treating all models equally. Bose et al. used “frequency of occurrence” for variable selection [21], which is essentially the most-votes method. In many other studies, researchers have listed the variable importance of all models but have not provided an overall measure of the relative influence of predictors [18,20,24,25,36]. Since different models perform quite differently in risk prediction and in variable selection, most-votes is a viable way to select the important variables, as most models selected these variables. Similarly, we may not want to treat all models equally in evaluating variable influence; i.e., unequal-weight weighted average of relative importance may be more reliable. In this study, we found that logistic regression–estimated weights outperformed simple average and most-votes in terms of prediction accuracy.

### Number of risk factors

The selection of the top 20 risk factors in this study was arbitrary. In our previous study of a multi-indicator approach for assessing malaria risks, we started with >200 variables, and the final GBM logistic model selected only 19 significant variables [43].

Similarly, in a study by Zheng et al. assessing monthly distributions of *Aedes albopictus* in China, they started with a similar >200 variables and the final model used only 17 variables [63]. Risk analyses by Solano-Villarreal et al. using boosted regression also ended up with 18 significant variables [64]. Several other disease risk analysis studies selected <20 predictors in their final models [65,66]. Practically, if we want to find the key risk factors, we should limit the number of candidate factors. In this context, we think the top 20 candidate predictors may be enough. In this study, each model selected 20 variables, there were about 50 candidate predictors in the ensembled model, but not all of them were equally important. In fact, in several models, such as SVM logistic and XGB tree, there were <5 key predictors based on their relative importance in the model, and the ensembled model only had 3–5 key predictors of habitats. Practically, selecting the top 20 important variables may be enough.

### Larval positive habitat identification

The accuracy of larval positive habitat predictions was low in this study. This is likely due to the selection of predictors at the begining. We used climatic and satellite observed environmental variables to predict the larval positive habitats. However, the presence of larvae in a habitat depends on two major factors: attractants for female breeding, and food and environment for larval development. Physical and chemical cues allow female mosquitoes to assess the suitability of potential larval habitats for breeding and hence influence the acceptance of oviposition sites [67–71]. Physical cues originate from vegetation (land cover type and density), water temperature, sunlight and texture of the substrate, and other biotic factors such as the existence of certain algae are crucial for larval development [72–76]. For example, Munga et al. found that land cover type affects *Anopheles* female oviposition [49]. Sumba et al. found that *An. gambiae* oviposition may be regulated by the daily light-dark cycle [77]. Eneh et al. found that water temperature also affects female oviposition site selection [78].

Factors such as vegetation cover, light-dark cycle and water temperature may be monitored by ground observations or satellite monitoring [73,74]. However, studies also found that certain biotic cues such as habitat microorganisms (e.g., bacteria) and volatile profiles (e.g., grass volatiles) affect female oviposition habitat selection [77–81]. These factors cannot be monitored through satellite monitoring or simple ground measurements. Biotic variables such as bacteria and grass types may vary from habitat to habitat and change over time. Therefore, it is entirely possible that while a certain aquatic habitat is suitable for female oviposition and larval development at this time, it will not be attractive for female oviposition and/or not suitable for larval development next time. Thus, predicting larval positive habitats is more difficult than predicting the aquatic habitats. In fact, very few studies have included larval positive habitat prediction [19].

The major limitation of this study is the selection of models for ensemble modeling. We don’t have a strategy for model selection, although data mining experts suggest selecting diverse and independent models for ensemble modeling [29,37]. We used diverse models in this study [17,25,26]. However, it was difficult to decide which models were independent or, more loosely, unrelated. We tried to select models that were not related to each other; we may also want to include other less-related models such as multiple adaptive regression splines, among others [25,26]. The second limitation is the selection of sites using field observations. Ideally, we should include diverse study sites. However, it is difficult to cover a wide variety of aquatic habitats with different ecological backgrounds due to the scope of scientific research; government census-style surveillance may include more diverse ecological settings. Including more diverse ecological areas may increase the accuracy of model predictions. The third limitation is the sample size in Kisii and Kisumu counties. Sample size was ∼35,000 in Kakamega and Vihiga counties, ∼6,000 in Kisii County, and ∼1,000 in Kisumu County; therefore, model results might be biased toward Kakamega and Vihiga counties, which have similar ecological conditions. This might be the cause of the low prediction probability in Kisii County and high prediction error in Kisumu County. Future modeling needs to consider balanced samples across different ecological settings.

In conclusion, this is the first study to provide a detailed framework for the process of multi-model ensemble modeling, including selection of models, estimation of model weights (statistically optimized), determination of key predictors (most-votes), evaluation of the relative influence of key predictors, risk mapping and prediction uncertainty assessment, including several key components of ensemble modeling that have not been addressed in previous studies. We found that elevation, geomorphon class and precipitation 2 months prior are the key malaria vector larval habitat predictors in western Kenya. We hope the modeling process we have proposed will be useful for other studies and that our predicted map of potential habitat availability will be helpful in assisting malaria vector larval source management planning.

## Acknowledgements

This study is funded by the National Institutes of Health (D43 TW001505, and U19 AI129326). The funders had no role in study design, data collection and analysis, decision to publish, or preparation of the manuscript.

## Supporting information

Figure S1. Flowchart of the modeling process

Figure S2. Accuracy of different ensembled models for the prediction of aquatic habitats (A) and larval positive habitats (B).

